# Crosstalk between RNA Pol II C-Terminal Domain Acetylation and Phosphorylation via RPRD Proteins

**DOI:** 10.1101/442491

**Authors:** Ibraheem Ali, Diego Garrido Ruiz, Zuyao Ni, Jeffrey R. Johnson, Heng Zhang, Pao-Chen Li, Mir Khalid, Ryan J. Conrad, Xinghua Guo, Jinrong Min, Jack Greenblatt, Matthew Jacobson, Nevan J. Krogan, Melanie Ott

## Abstract

Post-translational modifications of the RNA polymerase II C-terminal domain (CTD) coordinate the transcription cycle. Crosstalk between different modifications is poorly understood. Here, we show how acetylation of lysine residues at position 7 of characteristic heptad repeats (K7ac)— only found in higher eukaryotes—regulates phosphorylation of serines at position 5 (S5p), a conserved mark of polymerases initiating transcription. We identified the regulator of pre-mRNA domain-containing (RPRD) proteins as reader proteins of K7ac. K7ac enhanced CTD peptide binding to the CTD-interacting domain (CID) of RPRD1A and RPRD1B proteins in isothermal calorimetry and molecular modeling experiments. Deacetylase inhibitors increased K7ac- and decreased S5-phosphorylated polymerases, consistent with acetylation-dependent S5 dephosphorylation by an RPRD-associated S5 phosphatase. Consistent with this model, RPRD1B knockdown increased S5p, but enhanced K7ac, indicating RPRD proteins recruit K7 deacetylases, including HDAC1. We also report auto-regulatory crosstalk between K7ac and S5p via RPRD proteins and their interactions with acetyl- and phospho-eraser proteins.

## INTRODUCTION

The RNA polymerase II (Pol II) complex is highly conserved in all eukaryotic cells and responsible for production of most gene expression products (Buratowski, 2003; Eick and Geyer, 2013). RPB1, the largest subunit of the complex, contains the catalytic core of the complex and a unique regulatory region called the C-terminal domain (CTD). In eukaryotes, the CTD is composed of 20 or more repeats with a heptad consensus sequence, Y_1_S_2_P_3_T_4_S_5_P_6_S_7_, which is highly conserved from yeast to humans. In multicellular eukaryotes, the CTD is expanded and contains a varying number of non-consensus repeats, depending on the organism (Chapman et al., 2008). The 52 repeats of the mammalian CTD can be divided into 21 mostly consensus repeats proximal to the enzymatic core, and 31 non-consensus repeats distal from the core with less fidelity to the consensus. Divergence from the consensus sequence most commonly occurs at position 7, where the consensus serine is most commonly replaced with an asparagine (N), threonine (T), or lysine (K) (Eick and Geyer, 2013). The CTD is intrinsically disordered and functions as an interaction platform for accessory proteins required for transcription and transcription-associated RNA-processing events (Buratowski, 2009; Jasnovidova and Stefl, 2013).

The heptad repeats within the CTD are extensively and dynamically post-translationally modified at different times during the transcription cycle. Of the seven consensus CTD residues, five can be phosphorylated (Y1, S2, T4, S5, and S7), and the two remaining proline residues can undergo isomerization to *cis* or *trans* conformations (Heidemann et al., 2013). Serine-5 phosphorylation (S5p) and serine-2 phosphorylation (S2p) are the most thoroughly studied CTD modifications (Buratowski, 2009; Jasnovidova and Stefl, 2013). Serine-5 is phosphorylated by the cyclin-dependent kinase 7 (CDK7) subunit of general transcription factor TFIIH, is enriched at promoters, and decreases successively towards the 3′ end of genes (Brookes et al., 2012; Ebmeier et al., 2017). The phosphorylated serine-2 mark, placed by several kinases (CDK9, CDK12, CDK13 and BRD4), starts to accumulate downstream of transcription start sites and steadily increases towards the 3′ ends of genes, reflecting its critical role in productive polymerase elongation (Bartkowiak et al., 2010; Devaiah et al., 2012; Nechaev and Adelman, 2011). Similar to S5p, Serine-7 phosphorylation (S7p) is catalyzed by CDK7, is enriched near promoters and in gene bodies, and regulates the expression snRNA genes (Brookes et al., 2012; Egloff et al., 2012). Tyrosine-1 phosphorylation is enriched near promoters, and has been linked to enhancer and antisense transcription, as well as transcription termination (Descostes et al., 2014; Shah et al., 2018). Threonine-4 phosphorylation is enriched in coding regions and is required for cell viability and transcription elongation (Hintermair et al., 2012).

Post-translational modifications (PTMs) specifically found in non-consensus repeats include asymmetric dimethylation of a single arginine (R1810me2), conserved among some metazoa, that regulates transcription of small nuclear and nucleolar RNAs (Sims et al., 2011). In addition, lysine residues at position 7 of eight heptad repeats are acetylated by the acetyltransferase p300/CBP (KAT3A/B) (K7ac), and are also mono- and di-methylated by an as-yet-unknown methyltransferase (Dias et al., 2015; Schroder et al., 2013; Voss et al., 2015; Weinert et al., 2018). These lysine residues evolved in higher eukaryotes in the common ancestor of the metazoan lineage, and are highly conserved among vertebrates (Simonti et al., 2015). While lysine-7 mono- and di-methylation marks are found near promoters, K7ac is enriched in gene bodies (Dias et al., 2015). K7 residues are required for productive transcription elongation of immediate early genes in response to epidermal growth factor stimulation (Schroder et al., 2013). Importantly, K7ac marks are found at ∼80% of actively transcribed genes, with a peak in signal +500 bp downstream of the transcription start site (TSS), indicating that the modification could more broadly regulate the transition from transcription initiation to productive elongation (Schroder et al., 2013). In a genetic model in which all eight K7 residues were mutated to arginines (8KR), cells expressing 8KR RPB1 exhibited altered expression of genes relating to development, multicellularity and cell adhesion, underscoring a critical role of K7ac in the development of higher eukaryotes (Simonti et al., 2015).

Effector proteins interacting with differentially modified CTDs often contain a so-called CTD-interacting domain (CID), which is one of the best-studied CTD-binding modules and is conserved from yeast to humans (Ni et al., 2011). The mammalian <u>R</u>egulator of <u>P</u>re-m<u>R</u>NA <u>D</u>omain-containing (RPRD) proteins 1A, 1B and RPRD2 proteins are homologues of the yeast transcription termination factor *Rtt103*, and each contains a CID (Ni et al., 2011). *Rtt103* and RPRD CIDs bind CTD peptides carrying S2p, but not S5p; S7p and unmodified K7 residues reside at the edge of the CID binding cleft, and can be substituted without altering their binding affinity (Jasnovidova et al., 2017; Meinhart and Cramer, 2004; Ni et al., 2014). RPRD1A and RPRD1B are found in macromolecular complexes that associate with Pol II and transcription regulatory factors, including the S5-phosphatase RPAP2 (Liu et al., 2015; Morales et al., 2014; Ni et al., 2011; Ni et al., 2014; Patidar et al., 2016). RPRD1A, also called P15RS, regulates G1/S cell-cycle progression and suppresses Wnt and β-catenin signaling via interactions with the class I lysine deacetylase HDAC2 and transcription factor 4 (TCF4) (Jin et al., 2018; Liu et al., 2015; Liu et al., 2002; Wu et al., 2010). RPRD1B, also called CREPT, was identified in a mass spectrometry (MS)-based screen for mammalian Pol II-interacting proteins; it is upregulated in various cancers and regulates genome stability and transcription termination (Lu et al., 2012; Morales et al., 2014; Patidar et al., 2016; Zhang et al., 2018). Although the homology with *Rtt103* implies a conserved role in transcription termination and explains why the proteins are enriched in the 3′ ends of eukaryotic genes, an additional less well-defined role of RPRD proteins has emerged at the 5′ ends of genes in higher eukaryotes. This involves a mechanism to regulate genome stability through the resolution of R-Loops, which are DNA-RNA hybrids (Lu et al., 2012) as well as the recruitment of RPAP2 to initiating RNA Pol II (Ni et al., 2014).

In this study, we provide molecular insight into the role of RPRD proteins at the 5′ ends of genes and newly connect RPRD proteins with K7ac. We find that RPRD proteins via their CIDs specifically interact with K7ac, and that this interaction promotes S5-dephosphorylation at and beyond +500 bp downstream of the TSS. These data support a model in which vertebrates evolved specific crosstalk between S5p and K7ac to ensure precise transcription initiation dynamics and a timely transition to a productive elongation phase at a defined distance from the TSS.

## RESULTS

### Preferential Binding of RPRD Proteins to Acetylated RPB1

To identify proteins that interact with Pol II K7ac, we performed stable isotope labeling with amino acids in cell culture (SILAC). We overexpressed HA-tagged RPB1 proteins, either wild type (WT) or 8KR mutant, in HEK293T cells. The proteins also contained a known α-amanitin resistance mutation enabling propagation of successfully transfected cells in the presence of α-amanitin, which induces the degradation of endogenous Pol II (Bartolomei and Corden, 1987). After culture of cells in differential metabolic labeling medium, RPB1-containing complexes were purified via their HA tag and subjected to MS analysis (**Figure 1A, Supplemental table 1**). We found all members of the RPRD family preferentially bound to WT RBP1, including RPRD1A, RPRD1B, RPRD2 along with several of their interacting partners, such as RPAP2, RPAP3, MCM7 and RUVB1, that were identified by MS (Ni et al., 2011; Patidar et al., 2016) (**Figure 1B**).

**Figure 1:**
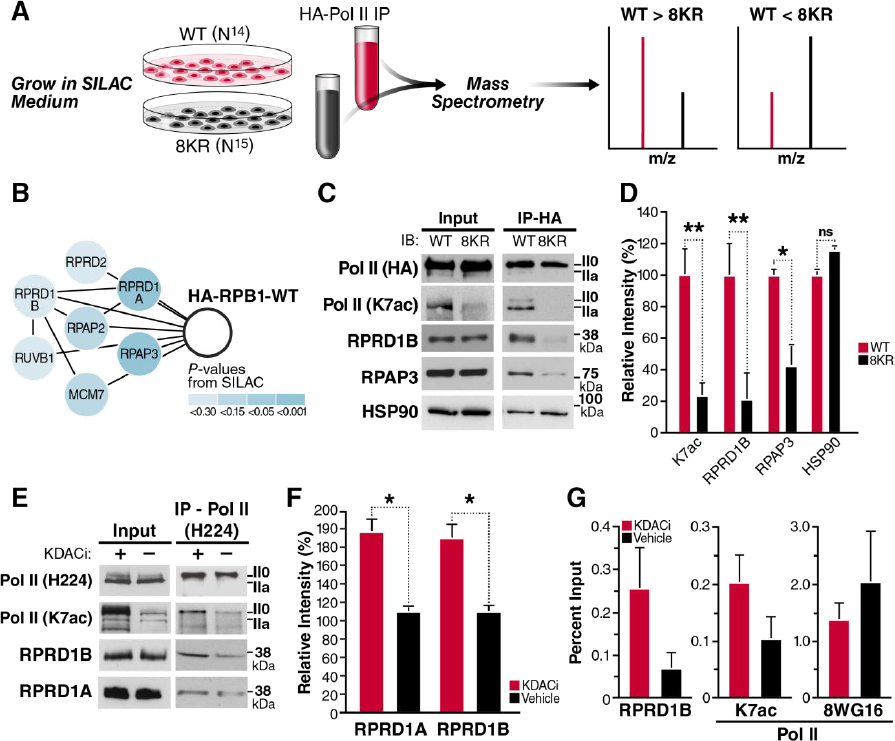
RPRD Proteins Interact with RPB1 in an Acetylation-Dependent Manner. (**A**) A SILAC-based mass spectrometry screen was used to identify factors that bind preferentially to WT Pol II, compared to the 8KR mutant. (**B**) Components of the RPRD1B/RPRD1A complex that were identified as WT interactors. (**C**) Hemagglutinin (HA) immunoprecipitation and western blotting with the indicated antibodies RPRD1B or from WT and 8KR cells. (**D**) Densitometry using ImageJ of at least three independent western blots from indicated antibodies after HA-immunoprecipitation. (**E**) NIH3T3 cells were treated with KDAC inhibitors (30 nM Panobinostat and 5 μM Nicotinamide) for 2 h. IP using an N-terminal H224 antibody from 500 μg of nucleoplasm pre-cleared with IgG and western blotting with the indicated antibodies. (**F**) Densitometry of RPRD1A and RPRD1B western blots produced from three independent Pol II Total (H224) IP elutions. (**G**) ChIP-qPCR on the *Leo1* gene at +33 nt downstream of TSS using the indicated antibodies from 4 independent chromatin preparations. Unmodified Pol II ChIP experiments were done with the 8WG16 antibody. Values are represented as percent of input with IgG subtracted. Error bars are SEM. *p < 0.5; **p < 0.01; ns, not significant for a one-tailed T test.

Using HA-immunopurification and western blotting, we confirmed enrichment of K7ac (p = 0.014) and RPRD1B (p=0.0098) in WT relative to 8KR samples, the most thoroughly studied of the RPRD family (Figure 1C). In addition, we confirmed that the RPRD-interacting protein RPAP3 (p = 0.049), binds preferentially WT RPB1 protein relative to the 8KR mutant. However, HSP90, another hit in the screen, showed no preferential interaction in follow-up studies (**Figure 1C, D**). The enrichment of endogenous RPRD1B and RPAP3 proteins after immunoprecipitation of WT, but not mutant, HA-RPB1 was consistent among three or more independent experiments and statistically significant (**Figure 1D**). We also tested the interaction of endogenous RPB1, RPRD1A and RPRD1B proteins in NIH3T3 cells treated with lysine deacetylase (KDAC) inhibitors. KDAC inhibitor treatment induced robust hyperacetylation of endogenous RPB1 in input material as tested with an antibody specific for K7ac (Schroder et al., 2013), but did not change unmodified Pol II protein levels as measured with the 8WG16 antibody, which correlate with total Pol II levels (Tsai et al., 2018; Vian et al., 2018) (**Figure 1E**). After pulldown of endogenous Pol II, more RPRD1A and RPRD1B proteins were recovered when cells were treated with KDAC inhibitors as compared to vehicle-treated cells, confirming positive regulation of the RPB1:RPRD interaction by acetylation **(Figure 1E, F)**.

Next, we tested *in vivo* recruitment of RPRD1B to a known target gene, *Leo1* (Ni et al., 2011). Using chromatin immunoprecipitation (ChIP) followed by quantitative PCR, we found RPRD1B recruitment to the *Leo1* promoter (+33 bp) consistently enhanced in NIH3T3 cells treated with KDAC inhibitors as compared to vehicle-treated cells (**Figure 1G).** Similar to what we observed by western blotting, K7 residues were hyperacetylated at the Leo1 promoter in response to KDAC inhibition in ChIP analysis with the K7ac-specific antibody. Importantly, Pol II occupancy as measured with the 8WG16 antibody did not increase under KDAC inhibition, confirming K7 hyperacetylation and enhanced RPRD1B recruitment in response to KDAC inhibition (**Figure 1G**).

### Direct Interaction of K7ac with RPRD CTD-Interacting Domains

To test whether K7ac modulates CTD-interacting domain (CID) binding to RPRD CIDs, we performed isothermal titration calorimetry (ITC) to measure the binding free energy between synthetic CTD peptides and purified CID domains from both RPRD1A and RPRD1B proteins. We generated CTD peptides spanning ∼3 heptad repeats (20 amino acids) with repeat 39 at the center. This region was chosen because it is acetylated and phosphorylated *in vivo* (Voss et al., 2015; Weinert et al., 2018), and contains multiple consecutive K7 residues (**Figure 2A**). Peptides were synthesized in an unmodified, acetylated (K7ac), or phosphorylated (S2p and S5p) state. S2p was included as a positive control as it enhances CTD:CID interactions, while S5p served as a negative control (Ni et al., 2014; Pineda et al., 2015). In addition, we combined S2p and S5p with K7ac to investigate potential combined effects. Compared to the unmodified CTD, binding of the RPRD CIDs to CTD peptides carrying K7ac had a significantly lower *K*_*d*_ (2.3-fold reduction for RPRD1A and 3.8-fold for RPRD1B), indicating enhanced binding (**Figures 2B–E**). S2p itself had a robust effect in enhancing CID binding, as previously observed, but combining K7ac with S2p further decreased the *K*_*d*_ by 2.8-fold and 4.2-fold, respectively. S5p-carrying peptides did not interact with CID proteins as expected, and adding K7ac did not further enhance binding (**Figures 2B–E**). These observations indicate that K7ac enhances the interaction of RPRD proteins with the Pol II CTD with and without additional S2p marks.

**Figure 2:**
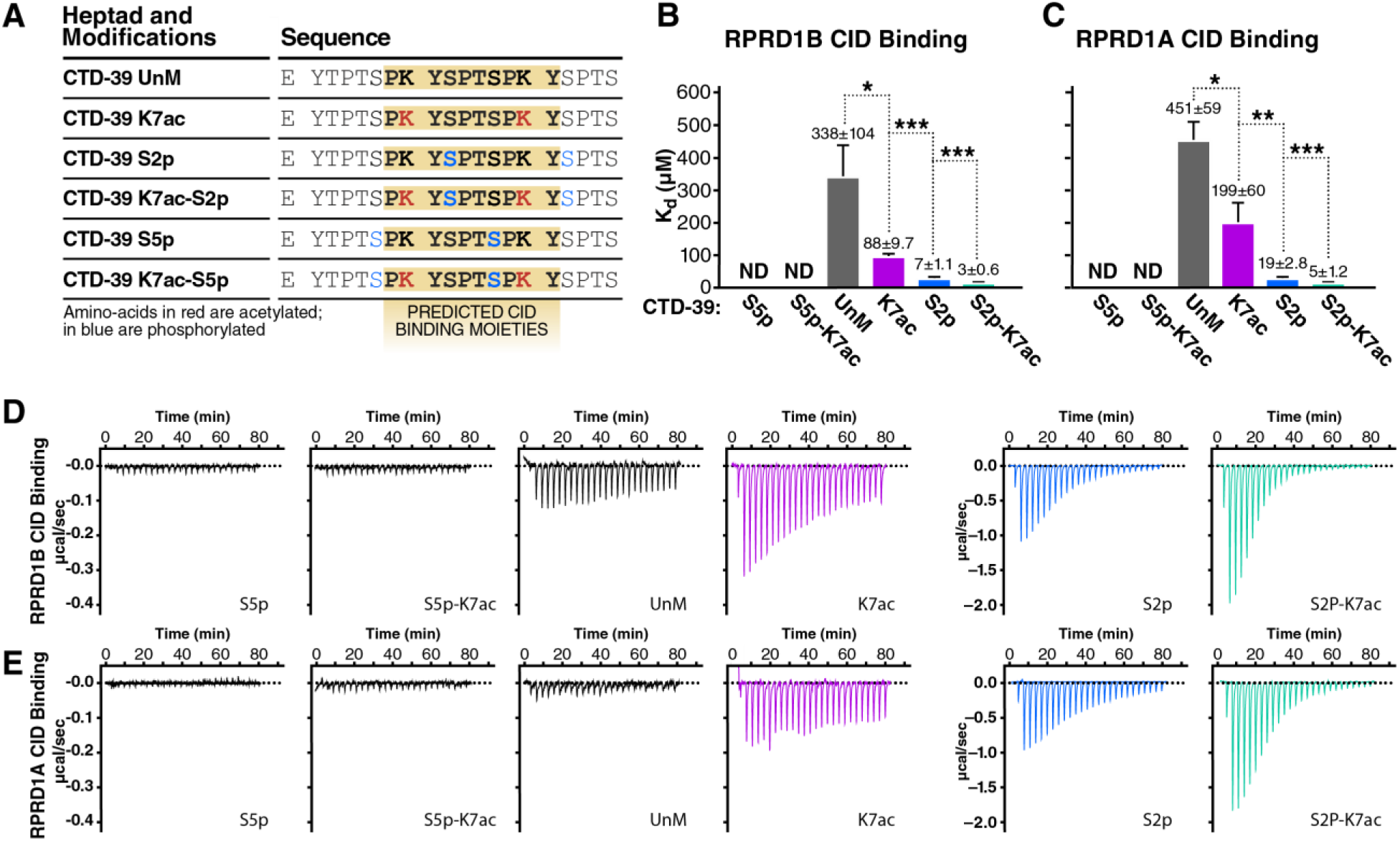
RPRD CID Domains Recognize Acetylated and Phosphorylated CTD Peptides. Isothermal titration calorimetry (ITC) experiments measuring *in-vitro* binding affinity between RPRD1A and RPRD1B CID domains and modified CTD peptides. **(A)** Table of CTD-39 peptides with indicated post-translational modifications and predicted RPRD CID binding moieties. **(B)** *K*_*d*_ values measured by ITC for RPRD1B CID and modified CTD-39 binding. **(C**) *K*_*d*_ values measured by ITC for RPRD1A CID and modified CTD-39 binding. **(D**) Representative ITC plots showing effect of K7ac on RPRD1B CID-CTD interactions for the indicated modifications. (**E)** Representative ITC plots showing effect of K7ac on RPRD1A CID-CTD interactions for the indicated modifications. * p < 0.05; ** p < 0.01; *** p < 0.005 using a one-tailed T test.

### Molecular Modeling of K7-Acetylated CTD Peptides with CID Domains

To better understand the mechanism for how K7ac stabilizes the binding of the CID and CTD peptides, we performed molecular modeling with published RPRD1B CID structures bound to CTD peptides [pdb: 4Q94 (dimer) and 4Q96 (tetramer)]. RPRD protein dimerization is believed to occur through coiled-coil domain interactions that are not present in these structures (Mei et al., 2014; Ni et al., 2014), which nonetheless dimerize and tetramerize by domain swapping. We proceeded with *in silico* analyses of both structures and searched for consistencies between them. Because phosphorylation and acetylation change the net charge of the peptide fragment, we first calculated the electrostatic potential of the CID structure to understand the charge distribution along the binding cleft (Dolinsky et al., 2004). We found that the recognition module within the CID in the dimeric and tetrameric structures has a positively charged binding pocket with numerous amide-containing residues within 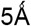 of the peptide (**Figures 3A–D, Supplemental Figure S1A–D**). Acetylation and thus charge neutralization of the CTD lysine side chain favors interaction with this binding pocket by reducing electrostatic repulsion with the positively charged RPRD domains.

**Figure 3:**
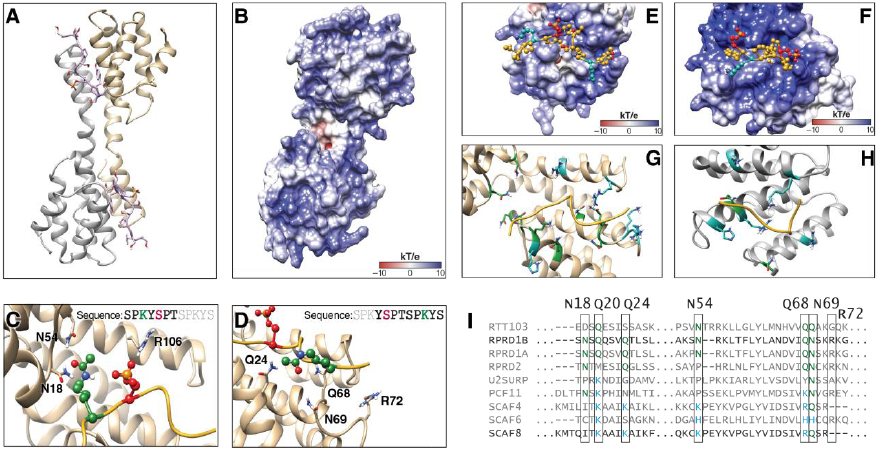
Molecular Modeling of K7 Acetylation and Interaction with CID Domains. **(A)** Dimer model from RPRD1B crystal structure (pdb:4Q94) containing two recognition modules and two peptides fragments of the CTD. **(B)**Electrostatic potential surface for RPRD1B CID. **(C)**Recognition elements around first K7ac and S2p in the CTD peptide fragment. **(D)**Recognition elements around second K7ac in the CTD peptide fragment. **(E)**CTD peptide fragment model from crystal structure superimposed with the electrostatic surface potential around the corresponding binding site. **(F)**CTD peptide fragment superimposed with the electrostatic potential surface around the corresponding SCAF8 CID binding site **(G)**RPRD1B CID residues found within 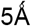 of peptide fragment. Blue: positively charged residues. Green: amide-containing residues. **(H)**SCAF8 CID residues found within 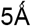 of peptide fragment. Blue: positively charged residues. Green: amide-containing residues. **(I)**Clustal Omega sequence alignment of human CID regions with yeast RTT103 as outgroup. Green: amide-containing residues. Blue: positively charged residues.

We also obtained 20-ns molecular dynamics trajectories to refine the models and identify residues within the CID that directly interact with S2p and K7ac and, thus, contribute to the recognition of these PTMs. Through these simulations, we reproduced the reported coordination between S2p and R106 (Ni et al., 2014), and observed that the two K7 residues in the acetylated state formed transient interactions with nearby CID residues (**Figure 3C, D**). In particular, the first acetylated K7 residue in the CTD peptide formed transient hydrogen bonds with two spatially proximal CID residues (N18 and N54). The second acetylated K7 residue interacted with CID residues at the other end of the binding cleft (Q24, Q68, N69 and R72) (**Figure 3C, D)**. Similar results were obtained in simulations of the tetrameric structure, which also showed coordination between S2p and R106 along with similar transient hydrogen-bonding between K7ac and several residues in the CID (N18, Q20, E92 and K96) (**Supplemental Figure S1E, F**). These results suggest that the acetylated lysine forms transient hydrogen bonding interactions that may contribute to binding stability, particularly with asparagine and glutamine that, like acetylated lysine, contain an amide group in the side chain.

Based on these observations, we hypothesized that RPRD1B evolved to specifically recognize K7ac through spatially proximal amide-containing side chains. Specific modes of recognition evolved to interact with different combinations of CTD modifications (Becker et al., 2008; Ni et al., 2014). For example, the SCAF8 protein recognizes doubly phosphorylated CTD peptides at S5 and S2 to regulate a putative role in pre-mRNA processing (Becker et al., 2008; Patturajan et al., 1998). Sequence alignment of CID domains reveals that RPRD proteins contain conserved asparagine and glutamine residues at key positions near the CID binding cleft. In contrast, the corresponding residues of SCAF family CID domains contain lysine, arginine or histidine residues (**Figure 3I**). We performed molecular modeling with acetylated non-consensus peptides that interact with the SCAF8 crystal structure (PDB: 3D9K). Interestingly, the SCAF8 CID binding interface contains a stronger net positive charge relative to RPRD1B with several lysine, arginine and histidine residues less than 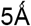 from the CTD peptide **(Figure 3E, F).** Only two amide-containing side chains were observed in the binding region (Q72, N120) compared with nine residues observed in RPRD1B **(Figure 3G, H)**. These data indicate K7 acetylation interacts with amide groups in stabilizing CID-CTD interactions in RPRD proteins, something not conserved across other CID containing proteins **(Figure 3I)**.

### Increased K7ac Correlates with Reduced S5p Downstream of Transcription Start Sites

RPDR1 proteins interact with RPAP2, the mammalian homolog of yeast *Rtr1* and a known S5 phosphatase (Ni et al., 2011). We tested the influence of K7ac on S5p levels by performing ChIP-seq with antibodies specific for K7ac, S5p (4H8) and unmodified (8WG16) Pol II in chromatin isolated from NIH3T3 cells treated with KDAC inhibitors. Average metagene profiles measured as reads per million, relative to input controls, were generated for all expressed genes (Ramirez et al., 2016) (**Figure 4A–C**). KDAC inhibition increased genome-wide Pol II-K7ac occupancy as expected, particularly ≥500 bp downstream of the TSS within the gene body and downstream thereof. Pol II occupancy as measured with the 8WG16 antibody was not markedly changed by KDAC inhibition (**Figure 4A, B**). Interestingly, S5p levels around the TSS remained unchanged, while beyond the ≥500 bp mark S5p levels decreased below the level of control cells, mirroring enhanced K7ac (**Figure 4C**). These data confirm that Kac and S5p levels are inversely correlated with a control point ≥500 bp downstream of the TSS.

**Figure 4:**
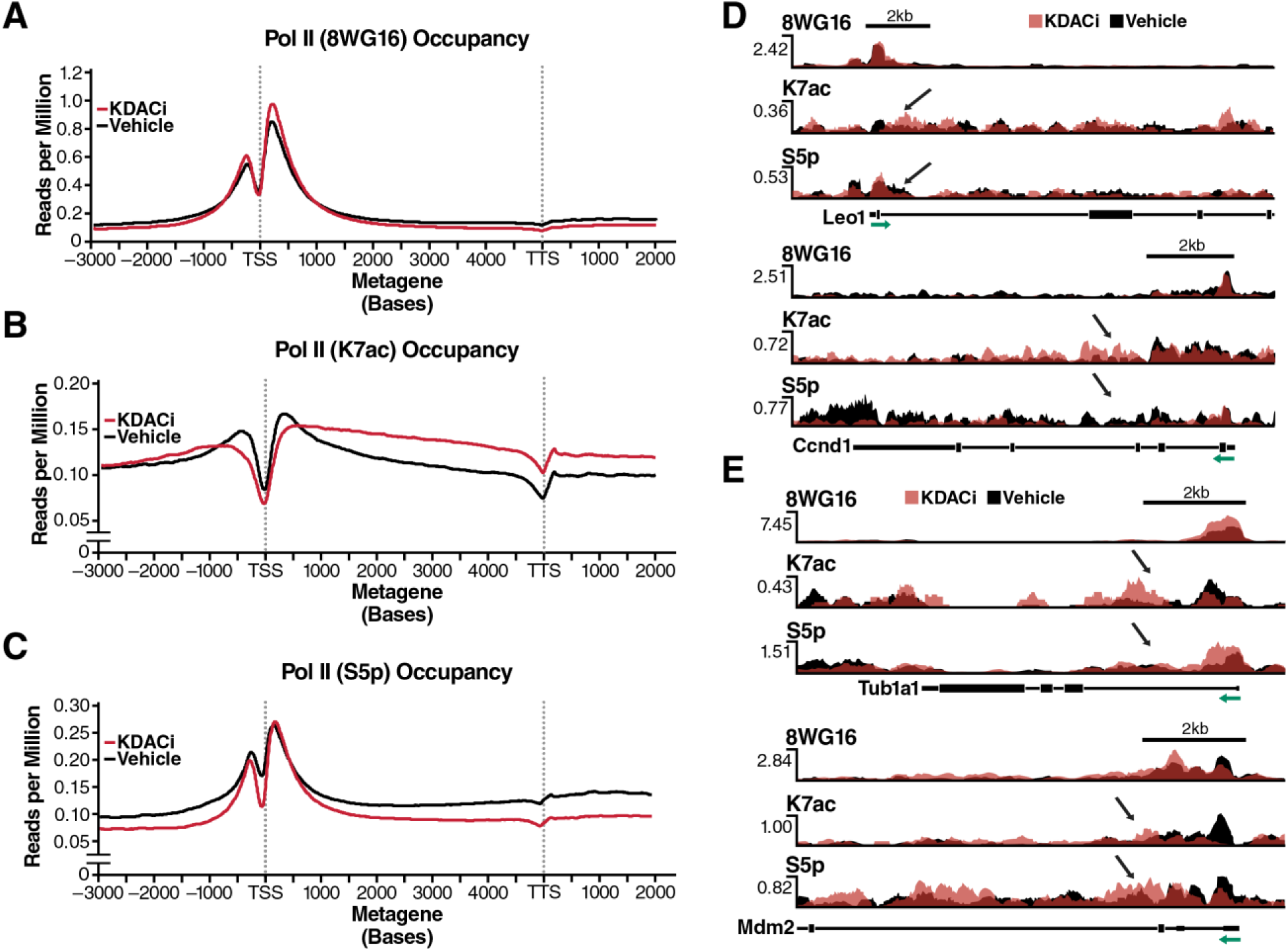
An Inverse Relationship between K7ac and Phosphorylation Is Induced upon KDAC Inhibition. **(A-C**) Metagene profiles generated from chromatin immunoprecipitation experiments in NIH3T3 cells with the indicated antibodies followed by deep-sequencing. Gene profiles are measured in reads per million normalized to input control **(D)** Single-gene validation of RNA Pol II PTMs measured as reads per million on selected RPRD1B occupied genes. S5p was measured using the RNA Pol II 4H8 antibody. **(E)** Occupancy profiles of RNA Pol II PTMs on control genes. Green arrows indicate the direction of transcription relative to the TSS of the depicted gene. Black arrows indicate the site of affected PTMs in response to HDACi.

A focused analysis of known target genes of RPRD1B, such as *Leo1* and *Cyclin D1* (Lu et al., 2012; Ni et al., 2011), showed corresponding profiles with lowered S5p and enhanced K7ac levels downstream of the TSS in response to KDAC inhibition (**Figure 4D**). But for ∼10% of actively expressed genes, including *Tub1a1* and *Mdm2*, S5p was not downregulated in response to KDAC inhibition despite strong upregulation of K7ac. These results indicate that these genes are possibly controlled by mechanisms other than the RPRD proteins (**Figure 4E**). Unfortunately, we could not examine the occupancy of RPRD proteins genome-wide as the available antibodies had an insufficient signal-to-noise ratio in ChIP-seq experiments (data not shown).

Monoclonal antibodies such as the 8WG16 and 4H8 antibodies that we use in this manuscript, are used to recognize CTD modification states by western botting and ChIP. Changes in signal, or absence of signal, can indicate either a physical absence of the modification, or masking of the epitope by other modifications (Heidemann et al., 2013). Using the peptides we used for our ITC experiments (**Figure 2A**), we tested by dot-blot our Pol II (K7ac) antibody specificity in the presence of S5p and S2p on the same repeat (**Supplemental Figure S2**). We observe that S5 or S2 phosphorylation decrease, but do not abolish the K7ac antibody recognition of acetylated CTD peptides. Furthermore, neither 4H8 nor 8WG16 antibodies are able to recognize the non-consensus CTD repeats tested (**Supplemental Figure S2**). These results indicate that the reduction we see in S5p is not likely due to masking of the 4H8 antibody epitope, and that we probably underestimate the signal of K7ac that we observe in our western blotting and ChIP-seq experiments.

### RPRD1B Knockdown Perturbs Both K7ac and S5p Marks Genome-Wide

Overexpression of RPRD proteins decreases S5p levels at the *Leo1* gene. This finding is consistent with a model in which RPRD proteins recruit the S5 phosphatase RPAP2 (Ni et al., 2011). We now performed the inverse experiment and knocked down RPRD1B in NIH3T3 cells using lentiviral shRNAs. A 50% knockdown efficiency was sufficient to induce global S5 hyperphosphorylation as observed by western blotting, indicating a critical role of RPRD1B in overall S5 dephosphorylation. Surprisingly, we also observed a consistent upregulation of K7ac levels in RPRD1B knockdown cells, suggesting a K7 deacetylase was recruited by RPRD proteins in addition to the S5 phosphatase **(Figure 5A)**. No change in Pol II levels as measured with the H-224 antibody was observed. Marked S5 hyperphosphorylation was also observed when the HA-8KR mutant RPB1 protein was immunoprecipitated from transfected NIH3T3 cells, confirming that K7 residues are required for proper S5 dephosphorylation **(Figure 5B).** In addition, we found endogenous HDAC1 protein co-immunoprecipitating with endogenous RPRD1B proteins in NIH3T3 cells, pointing to HDAC1 as a specific RPRD1-associated K7 deacetylase **(Figure 5C)**.

**Figure 5:**
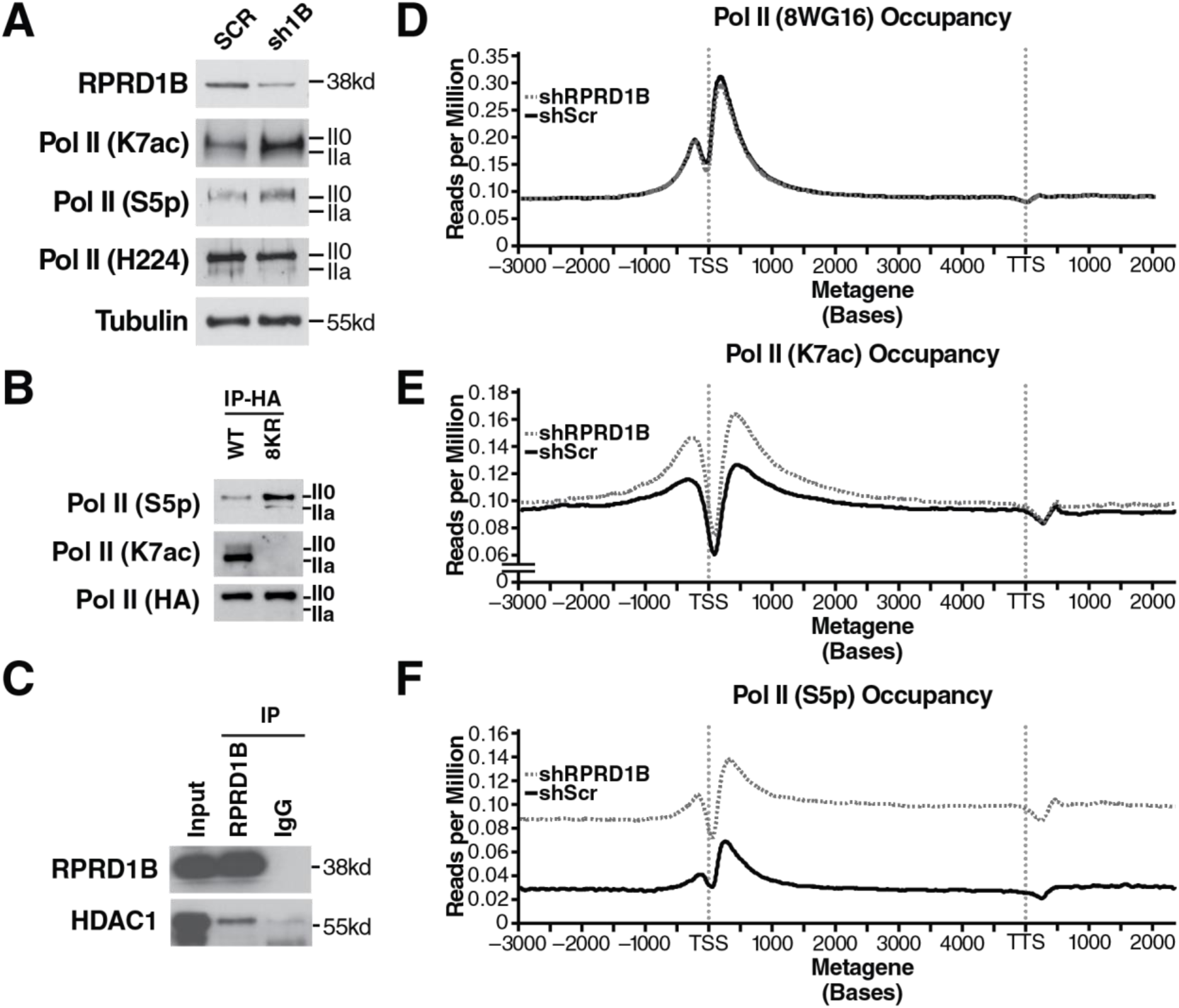
Increased S5p and K7ac Levels in Response to RPRD1B Knockdown. **(A)** Western blotting of indicated RNA Pol II modifications in NIH3T3 cells treated with scrambled shRNAs or those targeting RPRD1B. **(B)** Western blotting against S5p and K7ac from HA-immunopurified lysates containing WT or 8KR RNA Pol II. **(C)** RPRD1B immunoprecipitation from 293T cell lysates and western blotting against nuclear histone deacetylase HDAC1. **(D-F)** Metagene profiles of expressed genes in NIH3T3 cells. Gene profiles are measured in reads per million, relative to input control. Metagene profiles were generated from ChIP-seq data from NIH3T3 cells expressing either scrambled or RPRD1B targeted shRNAs treated with vehicle control. S5p is measured using the RNA Pol II 4H8 antibody. Profiles are representative of two independent experiments.

Next, we performed ChIP-seq in RPRD1B knockdown cells with antibodies against K7ac, S5p and unmodified Pol II. The most striking finding was the induction of a distinct TSS-proximal increase in K7 acetylation with minimal changes to 8WG16 Pol II occupancy (**Figure 5D and 5E**). This is consistent with the observation that K7ac was induced upon RPRD1B knockdown in western blot experiments (**Figure 5A**) and underscores a model where RPRD1B recruits a K7 deacetylase that counterbalances K7 acetylation within the first 500–1000 bp of transcribed genes. In conjunction with this, a dramatic increase in S5p upon RPRD1B knockdown (**Figure 5D, F**) fits well with the observation that reduced RPRD1B levels globally induce S5 hyperphosphorylation due to the lack of S5 phosphatase recruitment. Interestingly, K7ac enrichment was localized specifically to the TSS-proximal region, whereas S5p levels were higher globally than in cells treated with control shRNAs.

RNA-seq on RPRD1B knockdown cells identified 271 differentially expressed genes as compared to control shRNA-treated cells (**Supplemental Figure S3A, Table S2**). RPRD1B was among the most significantly downregulated genes with an mRNA knockdown efficiency of 41% (p = 0.0019; **Supplemental Figure S3D**), similar to what was observed for protein expression. Gene Ontology analysis on dysregulated genes indicated that RPRD1B knockdown induced changes in genes related to developmental processes, multicellular organismal development and cell adhesion, consistent with previous findings that complete knockout of the factor causes embryonic lethality (Morales et al., 2014) **(Supplemental Figure S3B–D).** This finding agrees with studies indicating that K7ac specifically evolved in higher eukaryotes and regulates developmental genes with significant enrichment for evolutionary origins in the early history of eukaryotes through early vertebrates (Schroder et al., 2013; Simonti et al., 2015). These data identify RPRD1B as a regulator of genes involved in multicellular organismal development and further support the model that RPRD proteins are relevant reader proteins of the K7ac mark in higher eukaryotes.

## DISCUSSION

In this study, we report a new molecular function for K7ac in decreasing S5p levels in the transition from transcription initiation to elongation. We show that K7ac enhances the recruitment of RPRD proteins to the initiated Pol II complex to facilitate S5 dephosphorylation; this occurs presumably via RPAP2, their interacting S5 phosphatase **(Figure 6A)**. Surprisingly, we found that lack of RPRD1B protein expression also increased K7ac levels, indicating that in addition to binding an S5 phosphatase, RPRD proteins recruit a K7 deacetylase that we identified as HDAC1. This provides a unique autoregulatory mechanism as binding to RPRD proteins to K7ac ultimately leads to the removal of the mark. Previous studies highlighted the importance of S2p in enhancing the interaction between the CTD and RPRD CID domains (Ni et al., 2014; Pineda et al., 2015), and so, the emergence of S2p downstream of K7ac may serve to maintain RPRD recruitment to complete S5 dephosphorylation during the early phase of transcription elongation (Ni et al., 2014). When levels of K7ac were perturbed by KDAC inhibitor treatment (**Figure 6B)** or RPRD1B knockdown (**Figure 6C**), recruitment of the S5 phosphatase was either enhanced, resulting in increased S5 dephosphorylation and lower S5p levels genome-wide, or recruitment was diminished, enhancing S5p levels, respectively. Therefore, these data support a model in which dynamics of K7 acetylation evolved to blunt the peak of S5 phosphorylation at a precise distance from the TSS in higher eukaryotes, likely facilitating the transition between transcription initiation and productive elongation.

**Figure 6:**
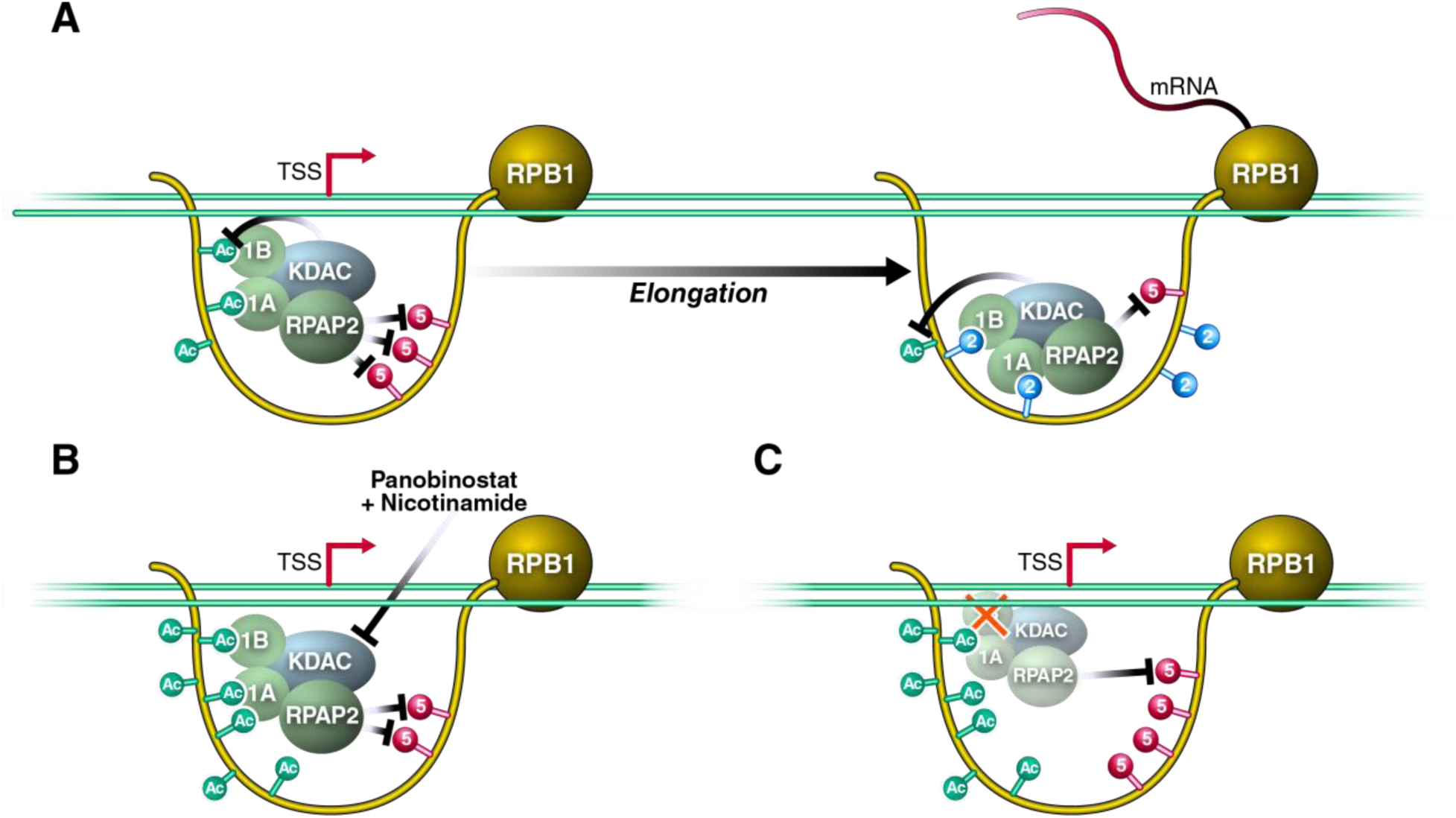
RPRD Proteins Are Recruited to the RPB1 CTD via Acetylation and Phosphorylation to Antagonize S5-Phosphorylation. (**A**) Model of RPRD complex reader and effector functions along modified residues within the CTD. 1B-RPRD1B 1A-RPRD1A, KDAC is a yet-to-be-identified K7 deacetylase. (**B**) KDAC inhibition induces K7 hyperacetylation and downregulation of S5-phosphorylation. (**C**) Knockdown of RPRD1B perturbs the recruitment of the complex to result in both S5 hyperphosphorylation and K7 hyperacetylation. See text for details.

We previously reported a transient enrichment in the occupancy of K7-acetylated Pol II located approximately +500 bp downstream of the TSS when we normalized K7ac peaks to unmodified Pol II occupancy on expressed genes (Schroder et al., 2013). This fits well with the self-limiting nature of K7ac where the modification recruits HDAC1 to support a negative feedback loop of K7ac via the RPRD reader proteins. Here we observed characteristic changes in S5p levels at and beyond the +500 bp mark that support the model that K7ac and S5p are regulated by RPRD proteins. In our current study, peaks proximal to the +500 bp mark behaved on average differently than peaks at or beyond the mark. There are several possible explanations. A) RPRD proteins are recruited specifically to the +500 bp location (at the peak of K7ac) and exert their effect on S5p and K7ac in this region and beyond. B) The balance between S5 phosphorylation and dephosphorylation immediately downstream of the TSS is shifted towards S5p, and efficient dephosphorylation only occurs after CDK7 levels are additionally lowered beyond the TSS (Ebmeier et al., 2017). C) The transition of Pol II into elongation and the occurrence of S2p at the transition point allow efficient RPRD association and S5 dephosphorylation and K7 deacetylation. We envision that multiple of the mechanisms could be in place to explain the observed changes.

An important finding of our study is that K7ac enhances the binding of the RPRD CIDs to CTD peptides. We show that the affinity of the CID:K7ac interaction is ∼90 μM, which lies in the range of acetyl-lysine interactions with bromodomains, the latter being the “classical” Kac recognition domain (Muller et al., 2011). Interestingly, the K7ac:CID interaction accommodates additional phosphorylation marks, such as S2p. This supports previous findings that the CID domain creates a positively charged “channel” in which the CTD peptide is bound, depending on its PTM status (Jasnovidova et al., 2017; Ni et al., 2014). Molecular modeling suggests that K7ac recognition occurs through electrostatic interactions and hydrogen bonding with various residues of the CID. The principles of specific recognition of phosphorylated amino acids have been well-studied and are consistent with the features we highlighted for S2p.

Although lysine acetylation has received less attention, our results suggest that side chains containing amide groups (N and Q) are important and form transient hydrogen bonds with the amide group of acetylated K7 residues. The amide groups in proteins, including those in the backbone and asparagine and glutamine side chains, tend to interact with each other, such as in secondary structure elements, asparagine capping the N-terminal end of alpha helices, and poly-glutamine repeats (Perutz et al., 1994). It is especially interesting that N18, N69 and Q24, which form hydrogen bonds with the acetylated lysine amide in the dimer or tetramer models, are conserved across RPRD proteins in mammals (Ni et al., 2014), but not in other CTD-binding proteins, such as SCAF8. Furthermore, we note that several CTD heptad repeats contain asparagine at position 7. We speculate that RPRD proteins may recognize such repeats, although this will require further experiments.

Serine-7 phosphorylation is the third-best studied PTM of the CTD. In mammals, S7p is enriched with S5p near promoters, but is uniquely stable in gene bodies (Descostes et al., 2014). Similar to K7ac, S7-phosphorylated residues are considered docking stations for RPAP2 that regulate S5-dephosphorylation and expression of snRNA genes (Egloff et al., 2012). Interestingly, S7p also enhances CTD:CID interaction consistent with the enhancement of electrostatic stability we describe here for K7ac (Egloff et al., 2012; Ni et al., 2014). Recent studies examined individual repeat PTMs *in vivo* and showed that CTD heptads are generally phosphorylated at one position per repeat (Schuller et al., 2016; Suh et al., 2016). This supports a model of dynamic movement along the 52 mammalian CTD repeats proposed here where the RPRD complex may start at distal non-consensus regions and work its way up to more proximal consensus regions to reach all S5p marks and allow maximal placement of S2p.

Another notable finding is the autoregulatory nature of K7ac. Although RPAP2 is a known S5 phosphatase, we showed that HDAC1 is a new RPRD1B-associated deacetylase. Previous studies with overexpressed proteins identified HDAC2 as a RPRD1A-associated deacetylase and, thus, another possible candidate for CTD regulation (Liu et al., 2015). However, using endogenous proteins, we found no HDAC2 associated with RPRD proteins, but this may be due to limited sensitivity of the antibodies used (data not shown). The two candidates, HDAC1 and HDAC2, are consistent with our previous observations that class I/II KDACs are involved in deacetylation of the hypophosphorylated form of Pol II during or after transcription initiation (Schroder et al., 2013). Notably, HDAC1 and HDAC2 often function together within the NuRD complex, and further studies will investigate how NuRD complex proteins could regulate K7ac and thus Pol II-dependent transcription (Basta and Rauchman, 2015; Xue et al., 1998).

Gene expression changes as a consequence of RPRD1B knockdown were moderate, but the cellular pathways altered in response to RPRD1B knockdown revealed a relevant list of genes. These were mainly involved in development of multicellular organisms and were strikingly similar to differentially regulated pathways found in WT and 8KR-Pol II expressing cells (Simonti et al., 2015). We have previously shown that K7ac evolution in higher eukaryotes presented a unique mode by which transcription elongation is regulated in mammals. We propose that the regulation of K7ac is linked to the now reported recruitment of RPRD proteins and the corresponding S5 dephosphorylation, a step tightly controlled in its dynamics in yeast. The question of why the need arose to control S5p with K7ac in multicellular organisms at a defined distance from the TSS remains unanswered but will be further examined. At this point, our data underscore a key role of controlled CTD PTM regulation at the transition from initiation to elongation important for the expression of developmentally relevant genes; they further demonstrate that this control depends on precise interactions with the RPRD complex, which performs reader and effector functions at a well-defined time during the transcription cycle.

## MATERIALS AND METHODS

### Antibodies and Reagents

See **Table S3** (Key Resource Table); Dynabeads Protein G (ThermoFisher, 10003D), Dynabeads Protein A (ThermoFisher, 10001D), bovine calf serum (Gemini, 100-506), 293T and NIH3T3 cells were from ATCC. Panobinostat (CAS 404950-80-7) and α-amanitin (CAS 23109-05-9) were purchased from Santa Cruz Biotechnology. All other chemicals and reagents were purchased from Sigma.

### Cell Fractionation and Immunoprecipitation

Cell fractionation was performed using the Dignam & Roeder method with minor modifications. 293T or NIH3T3 cells were pelleted and washed in cold DPBS. Pellets were resuspended in 5 volumes of DR Buffer A (10 mM HEPES-KOH, pH 7.9, 10 mM KCl, 1.5 mM MgCl_2_, 0.5 mM DTT, 1x HALT, 30 nM Panobinostat and 5 μM nicotinamide). Cells were Dounce homogenized with 10 strokes using a tight pestle (Wheaton), and cytoplasmic lysates were set aside or decanted. Nuclear pellets were resuspended in DR Buffer C (20 mM HEPES, 0.42 M NaCl, 1.5 mM MgCl_2_, 0.2 mM EDTA, 25% glycerol, 0.5 mM DTT, 1x HALT, 30 nM Panobinostat and 5 μM nicotinamide), sonicated using the Sonic Dismembrator 500 (ThermoFisher Scientific). 500 μg of nucleoplasm was precleared with normal IgG (Santa Cruz) conjugated to the appropriate beads, and immunoprecipitation was performed using anti HA-agarose beads (Sigma, A2095) or antibodies bound to Dynabeads. Immunoprecipitates were eluted either by boiling in 2x Laemmli buffer (agarose) or incubating in Elution Buffer (50 mM NaHCO_3_, 1% SDS) and adding 2x Laemmli buffer (Dynabeads).

### Lentiviral Transduction of RPRD1B shRNAs

VSV-G pseudotyped lentiviruses were produced to contain a puromycin resistance gene and a shRNA against RPRD1B (NM_027434.2-1003s21c1) or a scrambled control. Cells were transduced with 0.5 mL of unconcentrated virus and selected using 2 μg/mL puromycin for 1 week prior to experimentation.

### Chromatin Immunoprecipitation in NIH3T3 Cells

NIH3T3 cells were grown under normal conditions (10% BCS, 1x penicillin and streptomycin, 2 mM L-glutamine). We treated 6×10^7^ cells with a lysine deacetylase inhibitor cocktail (30 nM Panobinostat, 5 μM nicotinamide) or a vehicle control (DMSO, water) for 2 h. Cells were fixed with 1% formaldehyde for 15 min, thoroughly washed with DPBS, and resuspended in ChIP lysis buffer #1 (10 mM Tris, pH 7.4, 10 mM NaCl, 0.5% NP-40, 1x HALT, 30 nM Panobinostat and 5 μM nicotinamide). After sitting on ice for 10 min, cells were briefly vortexed, and the nuclei were pelleted. Nuclei were treated with MNase (NEB, M0247S) for 25 min at RT, pelleted and resuspended on ice in ChIP lysis buffer #2 (50 mM Tris HCl, pH 8.0, 10 mM EDTA, 0.5% SDS, 1x HALT, 30 nM Panobinostat and 5 μM nicotinamide). Chromatin was further sheared by sonication using the Sonic Dismembrator 500 (ThermoFisher Scientific) and preserved at −80°C until immunoprecipitation. 20–40 μg of chromatin was used for each IP with the antibody concentrations listed in **Table S3**. IPs were diluted into a final volume of 800 μL with ChIP Dilution buffer (167 mM NaCl, 16.7 mM Tris HCl, pH 8.0, 1.2 mM EDTA, 1.1% Triton X-100, 0.01% SDS) and left at 4°C overnight. IPs were washed then eluted in ChIP elution buffer (50 mM NaHCO_3_, 1% SDS) and decrosslinked at 65°C for 16 h. Samples were treated with RNAse A (Thermofisher, EN0531) for 20 min, and DNA was purified using the QIAquick PCR purification kit (Qiagen, 28106). Primer sequences are available upon request. For samples that were deep-sequenced, 2 ng of immunoprecipitated DNA from each reaction was used to create libraries using the Ovation Ultra-Low Library prep kit (Nugen, 0344-32), following manufacturer recommendations, and libraries were deep sequenced on the HIseq 4000 or NextSeq 500 using single-end 50 bp or single-end 75 bp sequencing, respectively.

### RNA Sequencing

RNA was prepared from 1×10^6^ NIH3T3 cells using the QIAgen RNeasy Plus Kit. Libraries were prepared with the Ovation Ultralow System V2 kit pn: 7102-32 / 0344-32, and libraries were deep sequenced on NextSeq 500 using paired-end 75 pb sequencing. RNA seq analysis was done using the Illumina RNAexpress application v 1.1.0.

### ChIP-seq Data Analysis

Barcodes were removed and sequences were trimmed using Skewer (Jiang et al., 2014). For each ChIP 50-60 million reads were aligned to the *Mus musculus* mm10 genome assembly using Bowtie with the –a -l 55 -n 2 -m 1 parameter (Langmead et al., 2009). Peaks were called and sequence pileups normalized to reads per million relative to input controls using MACS2 –B -SPMR -g mm -no-model -slocal 1000 (Zhang et al., 2008). TSS profiling was done using plotProfile on matrices generated with 10-bp bins using the computeMatrix function found in the Deeptools 2.2.3 build (Ramirez et al., 2016). Reproducibility of data was assessed by principal component analysis (Ramirez et al., 2016).

### Stable Isotope Labeling of Amino Acids in Culture (SILAC) of WT and 8KR Polymerases

SILAC labeling was performed according to the manual of SILAC Protein Quantitation Kit (LysC) –DMEM (Thermo Scientific cat. no. A33969). In brief, 293T cells stably expressing Pol-II-WT-HA and Pol-II-8KR-HA were grown in the light medium (L-Lysine-2HCl) or heavy medium (^13^C_6_L-Lysine-2HCl), respectively. After growing seven doubling times in the respective medium, incorporation efficiency of heavy L-lysine in 293T-Pol-II-8KR-HA cells was determined and the efficiency was more than 99%. To immunoprecipitate the HA proteins, 5 mg of total cell lysate from Pol-II-WT-HA and 5 mg of total lysate from Pol-II-8KR-HA cells in p300 lysis buffer were mixed together (total 500 μL), and 100 μL of HA agarose (Roche) were added. After overnight immunoprecipitation at 4°C, the HA-agarose was washed 4 times with 1 mL of cold p300 lysis buffer to remove non-specific binding proteins. The bound proteins were eluted twice by 100 μL of 0.1 M glycine, pH 2.5, after a 30-min incubation. Each elution was stored in separate tube. 10 μL of 1 M Tris-HCl, pH 8.0, was added into each elution to neutralize the pH. The quality of the elution was monitored by Protein Silver Staining (Pierce). Two elusions were combined and 50 μL out of the 200 μL combined elusions were analyzed by mass spectrometry (MS). Two independent biological repeats were performed.

### MS Analysis

Samples were analyzed on a Thermo Scientific LTQ Orbitrap Elite MS system equipped with an Easy-nLC 1000 HPLC and autosampler. Samples were injected onto a pre-column (2 cm x 100 μm I.D. packed with 5 μm of C18 particles) in 100% buffer A (0.1% formic acid in water) and separated by a 120-min reverse phase gradient from 5–30% buffer B (0.1% formic acid in 100% ACN) at a flow rate of 400 nL/min. The MS continuously collected spectra in a data-dependent manner, acquiring a full scan in the Orbitrap (at 120,000 resolution with an automatic gain control target of 1,000,000 and a maximum injection time of 100 ms), followed by collision-induced dissociation spectra for the 20 most abundant ions in the ion trap (with an automatic gain control target of 10,000, a maximum injection time of 10 ms, a normalized collision energy of 35.0, activation Q of 0.250, isolation width of 2.0 m/z, and an activation time of 10.0). Singly and unassigned charge states were rejected for data-dependent selection. Dynamic exclusion was enabled to data-dependent selection of ions with a repeat count of 1, a repeat duration of 20.0 s, an exclusion duration of 20.0 s, an exclusion list size of 500, and exclusion mass width of + or - 10.00 ppm.

Raw MS data were analyzed using the MaxQuant software package (version 1.2.5.8) (Cox and Mann, 2008). Data were matched to the SwissProt human proteins (downloaded from UniProt on 2/15/13, 20,259 protein sequence entries). MaxQuant was configured to generate and search against a reverse sequence database for false discovery rate calculations. Variable modifications were allowed for methionine oxidation and protein N-terminus acetylation. A fixed modification was indicated for cysteine carbamidomethylation. Full trypsin specificity was required. The first search was performed with a mass accuracy of +/-20 parts per million, and the main search was performed with a mass accuracy of +/-6 parts per million. A maximum of five modifications were allowed per peptide. A maximum of two missed cleavages were allowed. The maximum charge allowed was 7+. Individual peptide mass tolerances were allowed. For MS/MS matching, a mass tolerance of 0.5 Da was allowed and the top six peaks per 100 Da were analyzed. MS/MS matching was allowed for higher charge states, water and ammonia loss events. The data were filtered to obtain peptide, protein, and site-level false discovery rates of 0.01. The minimum peptide length was seven amino acids. Results were matched between runs with a time window of 2 min for technical duplicates.

### Isothermal Titration Calorimetry

RNA Pol II CTD peptides were purchased from Peptide 2.0 (Chantilly, VA). ITC experiments were performed as described (Ni et al., 2014)

### Molecular Modeling

Crystallographic structures were used for the dimer (pdb:4Q94) and tetramer (pdb:4Q96) models of RPRD1B and for SCAF8 (pdb: 3D9K). Electrostatic potential surfaces were calculated using an adaptive Poisson-Boltzmann solver (APBS) from the PDB2PQR server using the Amber force field and PROPKA to assign protonation states (Dolinsky et al., 2004). Amber’s LEaP program was used with the Amber ff14SB force field and the following force field modifications: phosaa10 (phosphates), ffptm (phosphorylated serines) and ALY.frcmod (acetyllysines). The TIP3P water model was used to solvate the system in a cubic periodic box, such that the closest distance between any atom in the system and the periodic boundary is 10Å. Net positive charge in the box was neutralized by adding counterions (Cl-) until neutrality. Energy minimization was performed in two steps: using harmonic restraints on the protein (10.0 kcal mol^−1^ Å^−2^) and an unrestrained minimization. For each minimization, we ran 1000 steps of steepest descent and 1000 steps of conjugate-gradient minimization at a constant volume with a non-bonded cutoff of 9Å. The equilibration was done in three steps. First, the system was heated from 0 to 300K with a restrained equilibration (10.0 kcal mol^−1^ Å^−2^) for 20 ps at constant volume with a non-bonded cutoff of 9Å, using the SHAKE algorithm to constrain bonds involving hydrogens and the Andersen thermostat. The second round of equilibration was performed by lowering the harmonic restraints (1.0 kcal mol^−1^ Å^−2^) on the system for 20 ps (other parameters identical). The third round was performed for 1 ns at constant pressure of 1.0 bar with non-bonded cutoff of 9Å at 300K with the Andersen thermostat. Simulations were performed without restraints using new velocities with random seeds at constant pressure of 1 bar with non-bonded cutoff distance of 9Å. 20-ns simulations were run with 2 fs timesteps per construct. Coordinates and energy were saved every picosecond (500 steps) (Case et al., 2005). Molecular graphics and analyses were performed with the UCSF Chimera package (Pettersen et al., 2004).

## Supporting information

SILAC Data Table

RNA-Seq Data Table

Resources Table

## ACKNOWLEDGEMENTS

We thank members of the Ott laboratory, J.J. Miranda, PhD, and Bassem Al Sady, PhD, for helpful discussions, reagents and expertise. Natasha Carli, PhD, and Jim McGuire from the Gladstone Genomics Core for library preparation and QCs for next-generation sequencing and for funding from the James B. Pendleton Charitable Trust. We are grateful for funding support from the NIH R01AI083139 (M.O.), P50-GM082250 (N.J.K.), and P01-CA177322 (J.R.J) UCSF Discovery Fellowship and American Society for Microbiology, Robert D. Watkins Graduate Research Fellowship (I.A.), and funding from the NSERC RGPIN-2016-06300 (J.M.). We thank John Carroll for graphics, Kathryn Claiborn for editing and Lauren Weiser for administrative support. Chimera is developed by the Resource for Biocomputing, Visualization, and Informatics at the University of California, San Francisco (supported by NIGMS P41-GM103311).

## AUTHOR CONTRIBUTIONS

I.A. designed the study, validated SILAC hits, conducted HDACi experiments, western blotting and quantification with assistance from M.K., performed ChIP-qPCR and ChIP-seq experiments and analyses, performed knockdown experiments, RNA-seq, and statistical analyses. P.C.L. and J.J. performed SILAC experiments and mass-spectrometric data analysis. Z.N., H.Z., and J.M. performed ITC experiments. D.G.R. performed molecular modeling experiments. R.J.C. supported ChIP-seq analysis. X.G., J.G., M.J., and N.K. supervised experiments. M.O. supervised the study design and data collection. I.A. and M.O. wrote the manuscript.

## SUPPLEMENTAL TABLES

**Table S1: Table of SILAC Mass-Spectrometry Data**

**Table S2: Genes Dysregulated by RPRD1B Knockdown**

**Table S3: Antibodies and Usage**

---

**Supplemental Figure S1:**
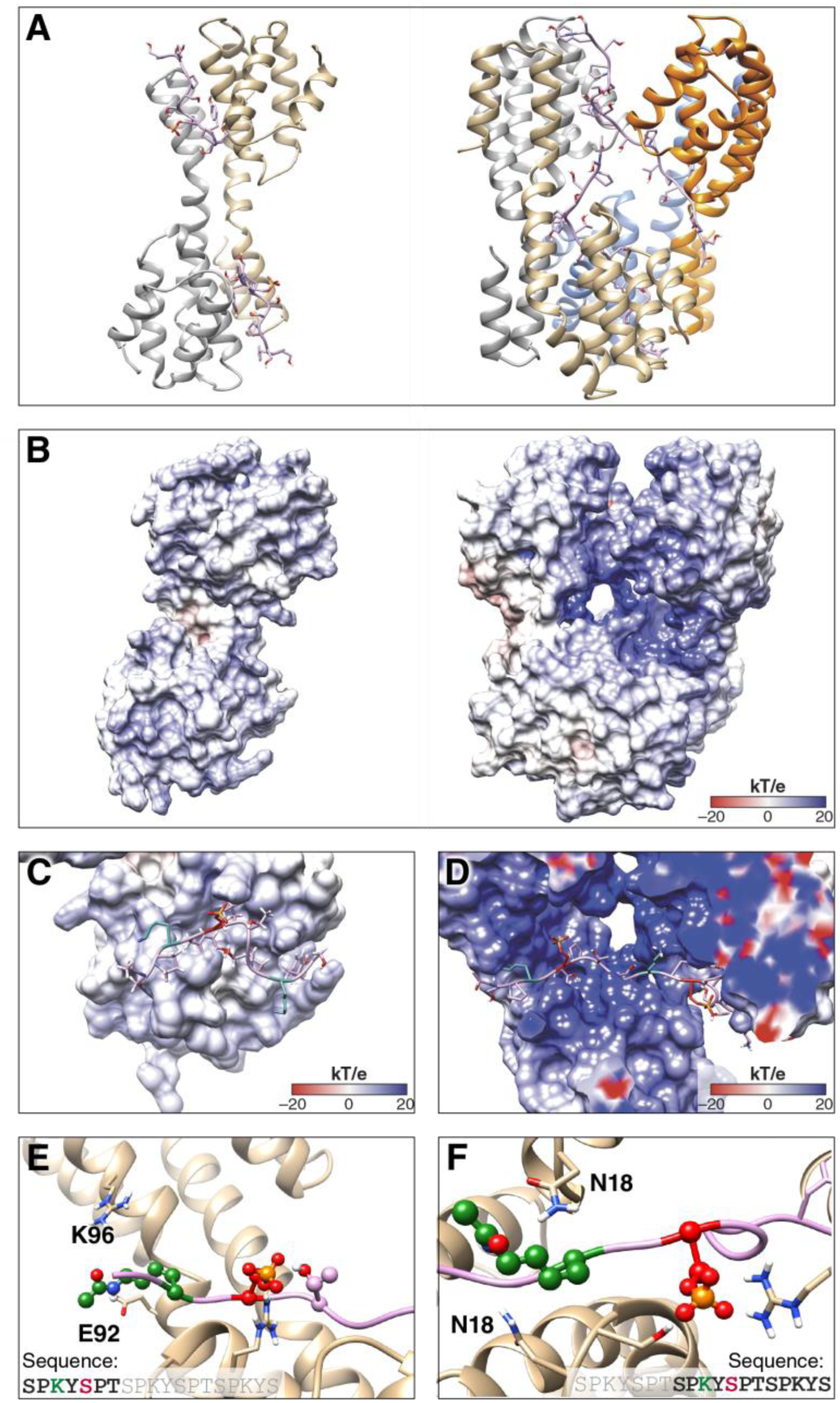
Dimer and Tetramer Models of RPRD1B:CTD Complex Share Similar Features. Comparison of dimeric and tetrameric models of RPRD1B:CTD complex. (**A**) Dimer (left) and tetramer (right) models from crystal structures. (**B**) Electrostatic potential surface for dimer (left) and tetramer (right) models. (**C**) Dimer model CTD peptide fragment superimposed with the electrostatic potential surface around the corresponding binding site. (**D**) Tetramer model CTD peptide fragment superimposed with the electrostatic potential surface around the corresponding binding site. (**E**) Recognition elements around first K7ac and first S2p in the CTD peptide fragment in the tetramer model. (**F**) Recognition elements around second K7ac and second S2p in the CTD peptide fragment in the tetramer model.

**Supplemental Figure S2:**
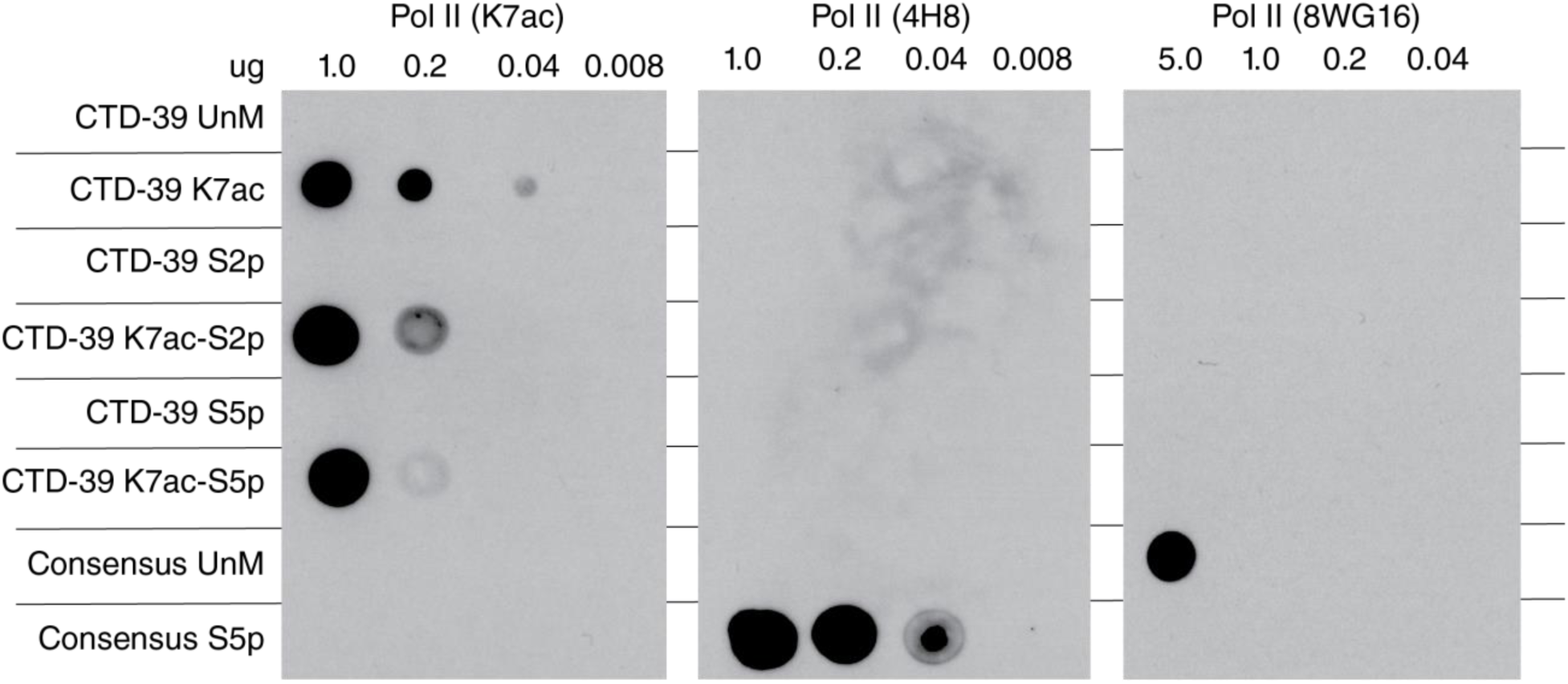
Antibody Specificity for Pol II (K7ac) and Recognition of Non-consensus CTD Heptads. CTD peptides centered around mammalian CTD repeat-39 (CTD-39) with the indicated modifications were blotted in a 5-fold dilution series to test the indicated anitbody specificity. UnM - Unmodified, K7ac - K7 acetylated, S2p - Serine 2 phosphorylated, S5p - Serine 5 phosphorylated. Consensus CTD peptides were used as controls for 4H8 and 8WG16 anitbodies.

**Supplemental Figure S3:**
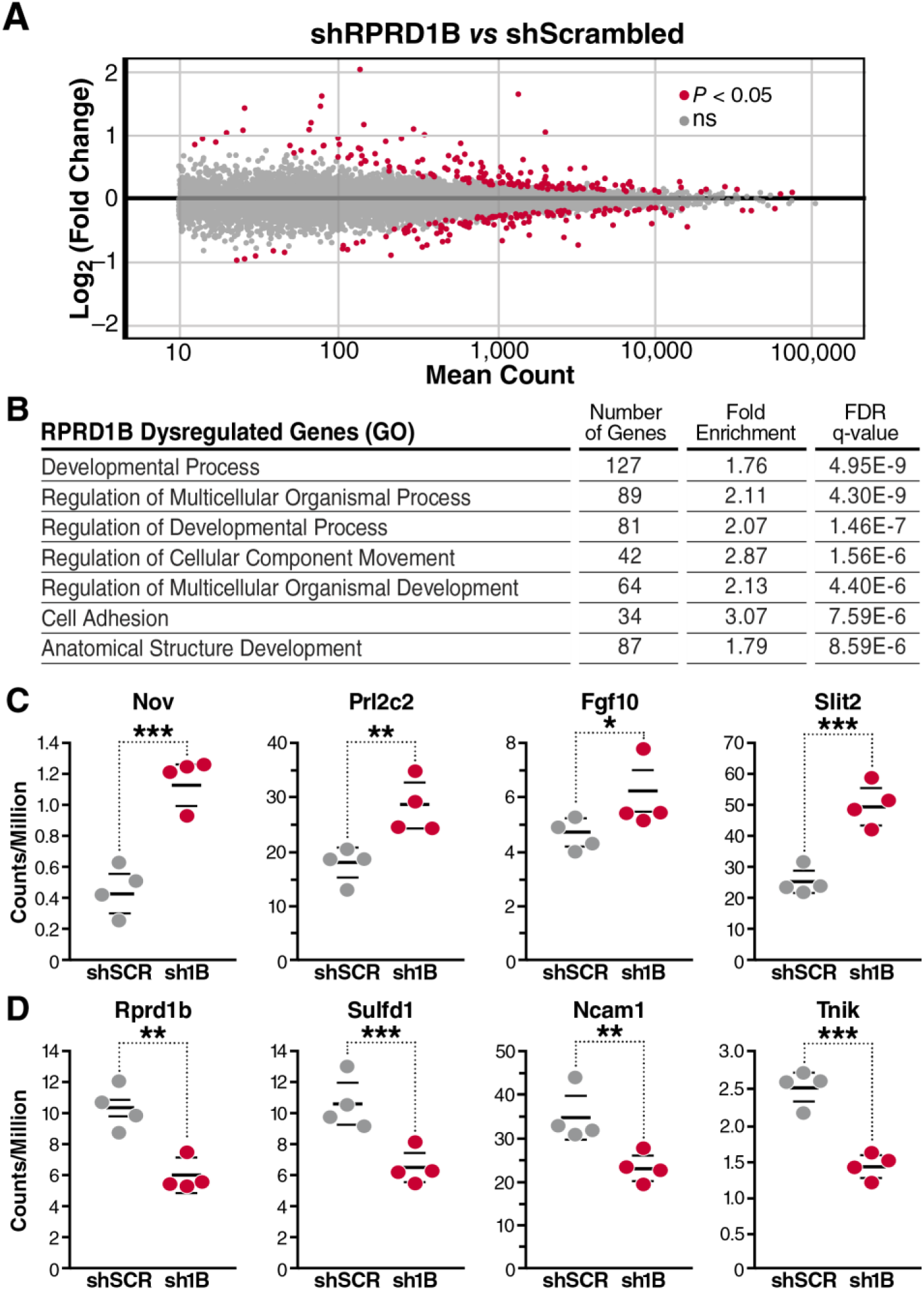
RPRD1B Knockdown Dysregulates Genes Relating to Development and Multicellularity. NIH3T3 cells were treated with shRNAs against RPRD1B (sh1B) or a scrambled sequence (shScr) and selected with puromycin for 1 week. (**A**) RNA-seq highlighting 271 significantly dysregulated genes in red. (**B**) Gene ontology analysis of genes significantly dysregulated in response to RPRD1B knockdown. (**C**) DESeq counts per million for selected upregulated genes associated with regulation of multicellular organismal process. (**D**) DESeq counts per million for selected downregulated genes in response to RPRD1B knockdown. * p < 0.05; ** p < 0.01; *** p < 0.005 using a one-tailed T test.

